# Devaluing memories of reward: A case for dopamine

**DOI:** 10.1101/2024.01.10.575106

**Authors:** B.R. Fry, N. Russell, V. Fex, B. Mo, N. Pence, J.A Beatty, F. P. Manfreddsson, B.A. Toth, C.R. Burgess, S. Gershman, A.W. Johnson

## Abstract

We describe a novel role for dopamine in devaluing sensory memories of reward. Mesencephalic dopamine cells activated during a mediated devaluation phase were later chemogenetically reactivated. This retrieval of the devalued reward memory elicited a reduction in the hedonic evaluation of sucrose reward. Through optogenetic and chemogenetic manipulations, we confirm dopamine cells are both sufficient and necessary for mediated devaluation, and retrieval of these memories reflected dopamine release in the nucleus accumbens. Consistent with our computational modelling data, our findings indicate a critical role for dopamine in encoding predictive representations of the sensory features of reinforcement. Overall, we illuminate the elaborate nature of reinforcement signals encoded by dopamine and suggest novel approaches to treating a host of psychobiological disorders.

## Main Text

Animals anticipate long-term future rewards in order to behave adaptively in the present. To achieve this, they can learn predictions of future reward directly from their experiences and update these estimates using reward prediction errors (RPE) ^1,2^. Traditional models of dopamine function posit that dopamine cells encode RPE ^3–5^. Although these studies have substantially increased understanding of dopamine function, other findings suggest dopamine neurons function in a more heterogenous manner during learning, as they fire in response to aversive cues^6^, novelty signals independent of value ^7^, and are necessary for encoding detailed features of the reward environment that are beyond the signal computed by RPE ^8–10^. With these findings in mind, more recent theories of dopamine function suggest these cells also encode a richer array of features from the environment ^11,12^. Consistent with these ideas, here we show a novel function for dopamine in the devaluation of detailed sensory features of reward through mediated learning.

### Using representation mediated learning to devalue reward

To reveal the potential vast array of features underlying dopamine-dependent learning and memory, we adapted a representation mediated learning approach, in which early on in training, a reward-paired conditioned stimulus (CS) can retrieve memories of food rewards so detailed in nature that animals sensorially experience the absent reward ^13,14^. This is presumed to reflect the capacity of the CS to activate perceptual processes within brain circuitry that are typically recruited by the presentation of the reward alone ^15^. Mice received Pavlovian training in which an auditory CS was paired either 16 (minimal) (Supplementary Fig. 1A) or 64 (extensive) (Supplementary Fig. 1E) times with 0.2M sucrose, followed by a memory devaluation phase in which the CS alone was presented and preceded an injection of the gastric malaise inducing agent, LiCl (see, Supplementary Methods). If the CS could retrieve detailed sensory memories of the previously paired but absent sucrose solution, we would expect the devaluation produced by LiCl to diminish the perceived palatability of the sucrose when it was reintroduced ^16^. To test whether mediated devaluation was achieved, we integrated quantitative analyses of rodent licking behavior, in which the temporal distribution of interlick intervals can be used to infer the perceived palatability of a consumed liquid reward ^17–21^. Despite the fact that the sucrose was never paired with LiCl, the devaluation of the detailed sensory features of sucrose reward was sufficient in minimally-trained mice to evoke a significant decrease in the perceived sweetness and palatability of the sucrose, compared to saline-control condition (Supplementary Fig. 1D).

### Activity-dependent labelling of ventral tegmental area cells to devalue memories of reward

To extend our examination of the neuronal basis of memory devaluation of sucrose reward, we labeled VTA cells in an activity-dependent manner during CS-evoked memory devaluation of sucrose reward, and reactivated these memories using chemogenetics. Transgenic cfos-htTA mice—in which the expression of tetracycline transcriptional activator (TRE) is directed to activated neurons by the c-fos promoter—received bilateral infusions into VTA of an adeno-associated virus (AAV) encoding the excitatory DREADD, hM3Dq, via pAAV-PTRE-tight-hM3Dq-mCherry (Fig. 1A). hM3Dq expression was only evident in mice that received doxycycline (Dox) withdrawal (Fig. 1B and 1C; Supplementary Figure 2). In mice that received activity-dependent labelling of DREADDs in VTA during aversion, treatment with clozapine-N-oxide (CNO) to activate hM3Dq-expressing cells and the putative devalued sucrose reward memory led to a bivalent effect over conditioned responding, such that pre-CS responses were elevated (Fig. 1D; p<0.05, d=0.45) and there was a tendency for CS responses to be reduced (Fig. 1E; p=0.1, d=0.39). Importantly, reactivation of these cells also attenuated both sucrose consumption (Fig. 1F; p=0.01, d=1.10), and its perceived palatability (Fig. 1G; p<0.05, d=2.23) but not the motivational to consume sucrose (Fig. 1H; p=0.1, d<0.5). These findings suggest that the VTA forms part of the neuronal circuitry underlying the encoding of detailed features of sucrose reward, such that they can be devalued and retrieved, leading to long-term changes in the evaluation of biologically meaningful rewarding events.

**Fig. 1.**
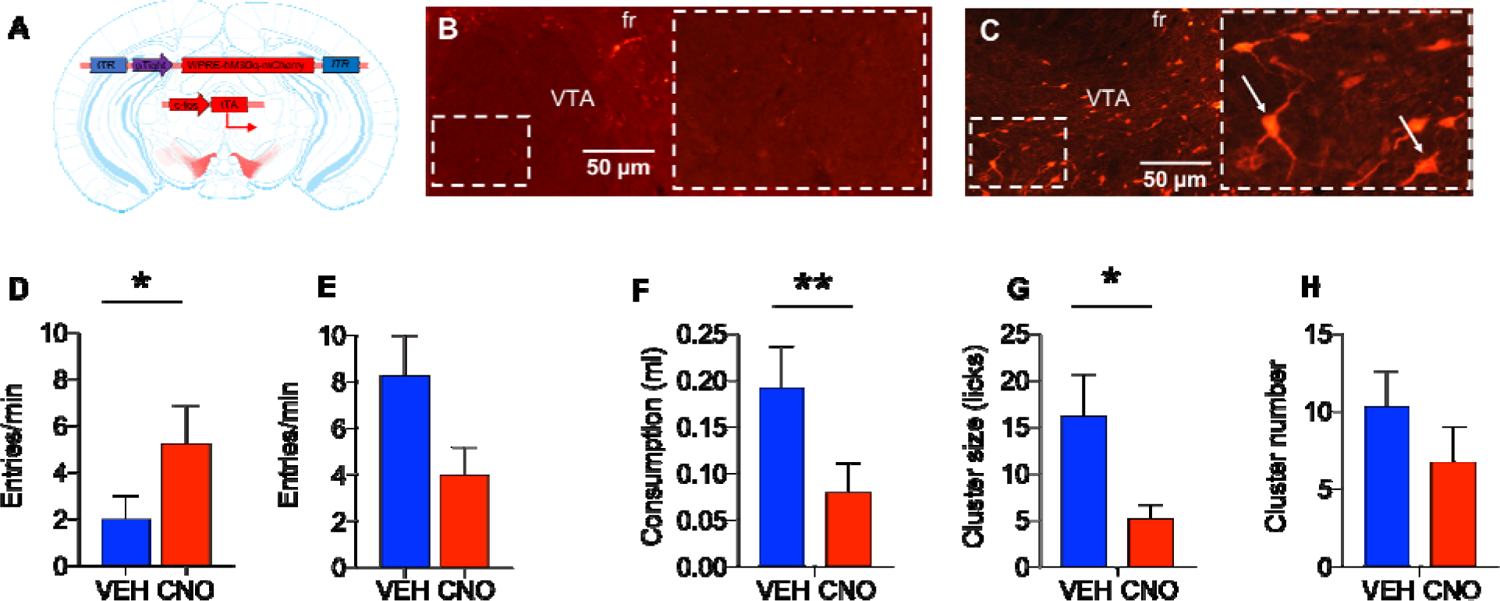
Mediated devaluation of sucrose reward via activity-dependent labelling and chemogenetic activation of ventral tegmental area cells. (A) cfos-htTA mice were bilaterally injected with pAAV-PTRE-tight-hM3Dq-mCherry. (B) cfos-htTA mice maintained on a diet containing Dox do not express hM3Dq-mCherry in the ventral tegmental area, whereas (C) while off Dox mice displayed extensive bilateral infection of hM3Dq-mCherry. Arrows indicate somatic expression of hM3Dq. (D-H) In the absence of Dox, cfos-htTA mice were trained to associate an auditory conditioned stimulus (CS) with 0.2M sucrose reward. During memory devaluation, the CS was used to reactivate a memory of the sucrose reward immediately followed by 0.6M LiCl injection. To label cells that were active during memory devaluation, cfos-htTA mice also received Dox injection and were maintained on a diet containing Dox for the remainder of the study. Two tests were then administered following vehicle and CNO intraperitoneal injections. During memory retrieval we examined whether the CS entered into associations with LiCl (CS test) and its capacity to mediate a devaluation to the sucrose reward (consumption). (D) Reactivation of devalued memories of sucrose enhanced pre-CS responding (E) but did not significantly influence CS responses. (F-H) Reactivation of devalued memories of sucrose reward significantly attenuated both (F) overall intake (G) and the palatability of sucrose reward, (H) but did not influence the motivation to consume it. Error bars indicate standard error of the mean (SEM). Abbreviations: fr = fasciculus retroflexus, VTA = ventral tegmental area. *p’s<0.05; **p=0.01.

### Ventral tegmental area dopamine cells are both sufficient and necessary for mediated devaluation of sucrose reward

Given this importance of the VTA in mediated devaluation of sucrose reward, we next examined whether stimulation of dopamine cells [the major neurotransmitter in the VTA^22^] would further enhance memory devaluation. Mice expressing Cre recombinase under the control of the tyrosine-hydroxylase (TH) promoter received unilateral injections of a Cre-dependent ChR2, (AAV5-Ef1α-DIO-ChR2-eYFP) or control eYFP (AAV5-Ef1α-DIO-eYFP) (Fig. 2A-D and Supplementary Fig. 3) along with an optical fiber cannula implanted slightly dorsal to the injection site. This approach allows temporally discrete and precise stimulation of ChR2-infected cells, resulting in depolarization in the presence of 473 nm optical stimuli ^23^ (Supplementary Fig. 4). VTA dopamine cell stimulation was timed to coincide with CS presentation during the aversion phase, with the prediction that this stimulation would promote further access to dopamine-dependent sensory features of reinforcement typically generated by the CS alone.

**Fig. 2.**
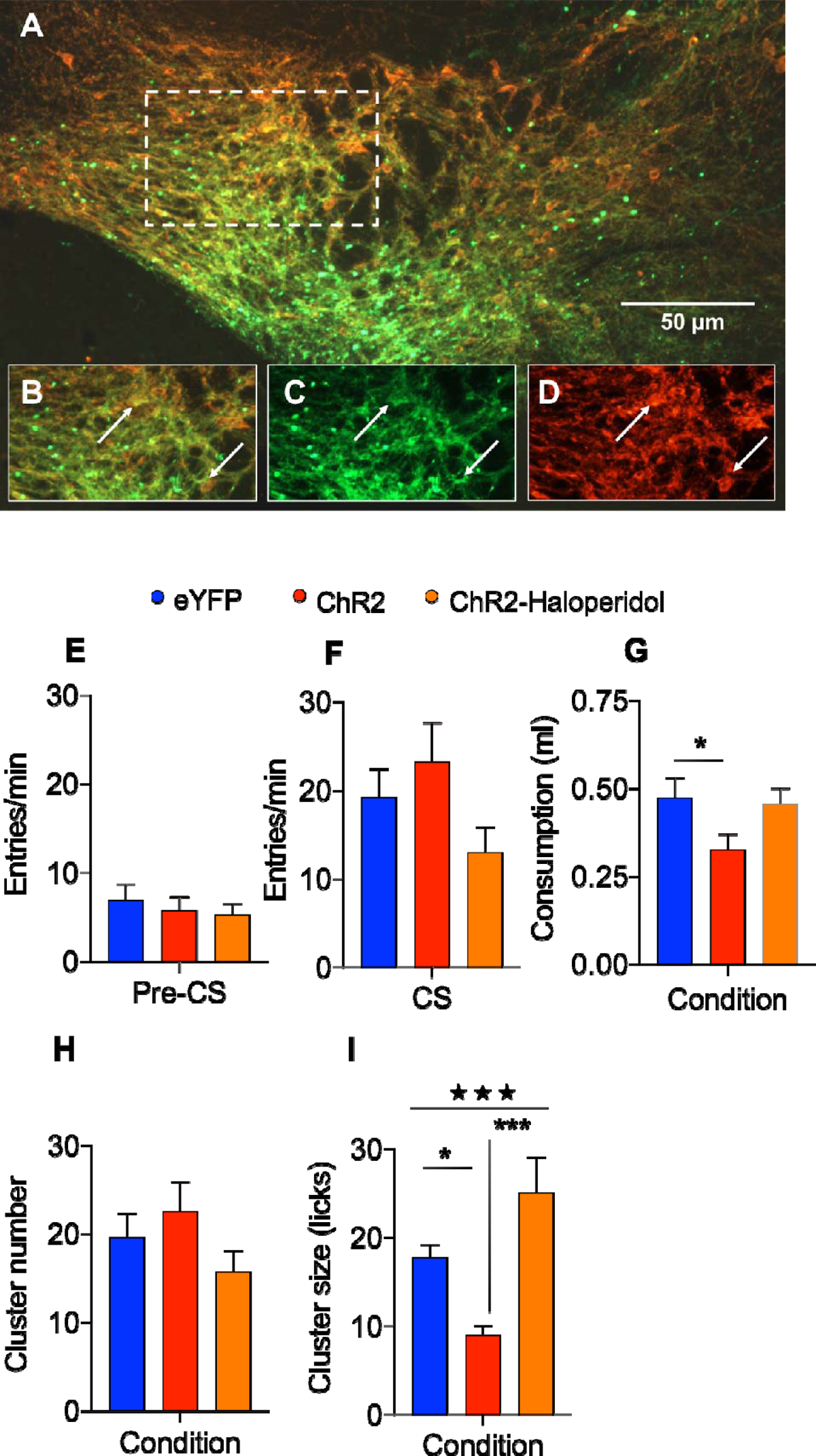
Optogenetic stimulation of ventral tegmental area dopamine cells enhances mediated devaluation of sucrose reward. **(A,B)** Representative photomicrographs displaying colocalization of ChR2 expression (green) in tyrosine hydroxylase (TH) positive neurons (red). Separate photomicrographs for **(C)** ChR2 and (**D)** TH. Arrows indicate somatic expression of ChR2 and TH. **(E)** pre-CS and **(F)** CS elicited food cup entries were comparable across all groups. **(G)** Optogenetic stimulation of VTA dopamine neurons enhanced mediated devaluation leading to a decrease in overall sucrose intake in ChR2 mice. **(H)** Cluster number was comparable across all group. **(I)** Relative to the mediated devaluation expressed by eYFP controls (blue bars), stimulation of VTA dopamine cells in ChR2 group (red bars) during aversion subsequently enhanced devaluation in the palatability of sucrose reward. This augmented effect was partly dependent on intact D2R signaling as peripheral treatment with haloperidol prior to optogenetic stimulation rescued performance to the level of eYFP in ChR2-Haloperidol group (orange bars). 111111 Overall group effect, p<0.0001. Post-hoc Bonferroni group differences, **\*\*\*** p<0.0001, **\*** p<0.05.

Accordingly, this would enhance the strength of the retrieved reward memory such that mice would express more robust mediated devaluation when the sucrose was reintroduced. Consistent with the idea that auditory stimuli do not reliably enter into associations with gastric malaise ^24^, no evidence was observed to indicate that the CS itself became devalued through direct pairing with LiCl, nor that optogenetic stimulation influences CS processing (Fig. 2F). However, VTA dopamine stimulation enhanced mediated devaluation as revealed by a significant reduction in overall intake between eYFP and ChR2 mice (Fig. 2G; p<0.05, d=1.0). The analysis of licking microstructure provided insight into the nature of this effect and indicated that willingness to initiate consumption of the sucrose solution was unaffected (Fig. 2H; p’s>0.26), whereas mediated devaluation for sucrose reward led to dramatic group differences (p<0.001) due to a reduction in the reward palatability of the sucrose solution between eYFP and ChR2 mice (Fig. 2I; p<0.05, d=2.39). This augmented mediated devaluation effect that followed VTA dopamine stimulation was dependent on intact signaling of the dopamine D2 receptor, as mice that received haloperidol (a D2 receptor antagonist) treatment prior to optogenetic stimulation displayed a pattern of licking behavior more similar to eYFP controls (Fig. 2G and 2I; p’s>0.09, d’s<0.55) and significantly different from ChR2 mice (Fig. 2I; p<0.0001, d=1.18). Furthermore, in a naïve group of ChR2 mice trained with 64 CS-US trials, which limits the expression of mediated devaluation (Supplementary Fig. 1G & 1H) ^15,25^, optogenetic stimulation during aversion was nevertheless capable of eliciting a significant decrease in the overall intake (Supplementary Fig. 5C; p=0.01, d=1.47) and perceived palatability of the sucrose (Supplementary Fig. 5E; p=0.01, d=1.45), compared to eYFP mice trained with the more extensive Pavlovian conditioning design.

To examine whether dopamine cell activity is necessary for mediated devaluation, TH-Cre mice received bilateral injections of a Cre-dependent inhibitory DREADD virus (AAV8-hSyn-DIO-hM4D(Gi)-mCherry) or control eYFP into VTA (Fig. 3A-D and Supplementary Fig. 6). All hM4Di-treated mice underwent mediated devaluation via CS-LiCl pairing; however, during this stage, for half the mice VTA dopamine cells were inactivated (hM4Di-CNO), whereas the remaining mice did not receive perturbations to these cells (hM4Di-VEH). The performance of these mice was compared to an eYFP control group that did not undergo mediated devaluation (eYFP) as they received saline rather than LiCl during the aversion phase. Subsequently, during the CS test, there was no evidence that the CS entered into an association with LiCl nor that inactivation of VTA dopamine cells during memory devaluation influenced pre-CS or CS responding (Figs. 3E & 3F). On the other hand, hM4Di-treated mice that did not undergo dopamine cell inactivation (i.e., hM4Di-VEH) displayed mediated devaluation for sucrose reward as evidenced by reduced overall intake (Fig. 3G; p<0.05, d>0.8), which did not impact the motivation to engage in sucrose intake (Fig. 3H; p=0.57, d<0.25) but reflected significant attenuation of reward palatability relative to eYFP mice that did not undergo mediated devaluation (Fig. 3I; p<0.01, d=3.08). Moreover, this refined measure of stimulus palatability also revealed an attenuated mediated devaluation of sucrose in hM4Di mice treated with CNO relative to those hM4Di mice that received vehicle prior to aversion (Fig. 3I; p<0.05, d=0.99). Further, inactivation of VTA dopamine neurons led to responding that did not statistically differ from eYFP mice that did not undergo memory devaluation (Fig. 3I; p=0.08). These findings confirm that in the absence of VTA dopamine cell activity, devaluation of the sweet-tasting features of sucrose reward failed to occur.

**Fig. 3.**
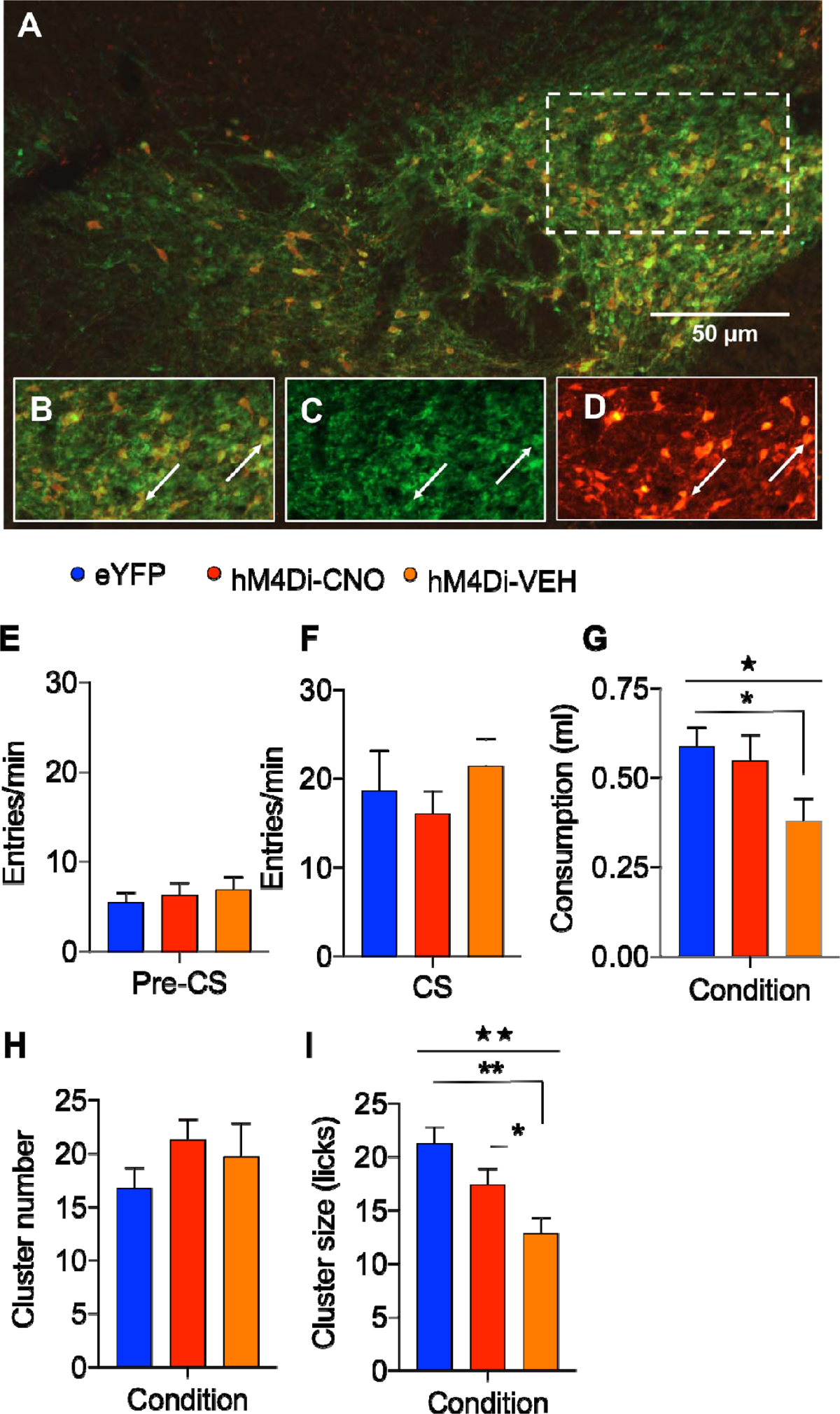
Chemogenetic inhibition of ventral tegmental area dopamine cells disrupts mediated devaluation of sucrose reward. **(A-D)** Representative photomicrographs of Cre-dependent hM4Di (red) in tyrosine hydroxylase (TH) positive neurons (green). **(E)** pre-CS and **(F)** CS elicited food cup entries were comparable irrespective of chemogenetic manipulations or whether mice received saline (eYFP-CNO) or LiCl (hM4Di-CNO, hM4Di-VEH) during aversion. **(G)** Chemogenetic inhibition of VTA dopamine cells (hM4Di-CNO) disrupted mediated devaluation relative to vehicle treated condition (hM4Di-VEH) and rescued performance to level of eYFP control mice that received saline during aversion phase (eYFP). **(H)** Inactivation of VTA dopamine cells (hM4Di-CNO) did not impact motivation to intimate consumption (cluster number); however, **(I)** only mice that underwent mediated devaluation in the absence of chemogenetic interrogation (i.e., hM4Di-VEH) displayed a reduction in palatability of the sucrose reward. By contrast, DREADD inactivation rescued performance in hM4Di-CNO group to the level of eYFP controls that were previously treated with CS-saline pairings (eYFP). Overall group effect, ★★ p’s<0.001. Post-hoc group differences, **\*\*** p<0.001, *p’s<0.05.

### Tracking physiological nucleus accumbens dopamine responses that underlie the retrieval of mediated devaluation of reward

One of the caveats of optogenetic and chemogenetic approaches is that they are unlikely to mimic endogenous dopamine activity. One of the major outputs of ventral mesencephalic dopamine cells is the nucleus accumens (NAc), a circuit critically implicated in dopamine-dependent RPE ^26^, incentive salience ^27^and sensorimotor activational processes^28^. Thus, to address whether VTA dopamine cell circuitry normally encodes detailed reinforcement signals consistent with a role in mediated devaluation, we used fiber photometry to dynamically track local dopamine release in the NAc. Mice received an AAV for the engineered dopamine receptor (dLight 1.1) into the NAc ^29^ (Fig. 4A). This fluorescent biosensor permits recording of GFP upon receptor binding of dopamine, producing fast observable responses to local dopamine release (Fig. 4B). Mice received Pavlovian training with two discriminable auditory CSs paired with the two distinct flavored pellets. This approach permitted a rigorous within-subject control for photometry to compare dLight activity. During the final stages of training, NAc dopamine responses displayed an expected increase in activity in response to the CSs and food USs ^26,30^, and importantly prior to aversion there were no differences in stimulus responses (Supplementary Fig. 7). When mice were tested to examine whether previously pairing the CS with LiCl altered its encoding, findings from the CS test (Fig. 4C) revealed comparable dLight activity in the NAc for each CS, irrespective of whether it was paired with LiCl or saline during aversion, which is consistent with the previously reported behavioral findings (Fig. 2F and 3F) indicating that CS processing was unaffected by mediated devaluation. Strikingly, however, during the consumption test we observed significant increases in dLight in the NAc as mice consumed the food pellet whose memory had been devalued by pairing its CS associate with LiCl (Fig. 4D; p<0.01, d=2.12).

**Fig. 4.**
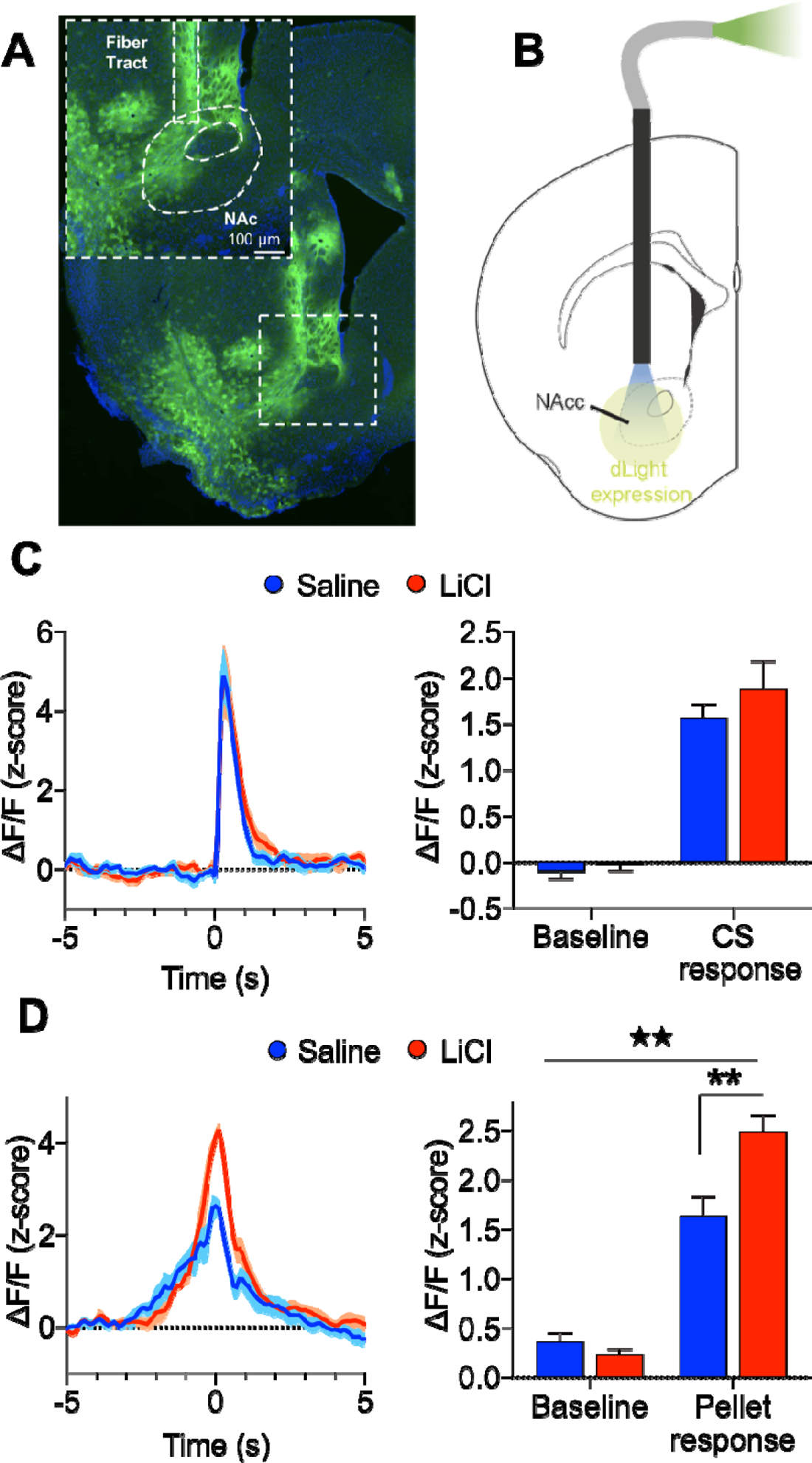
Dopamine release in nucleus accumbens is modulated by reward cue presentation and retrieval of mediated devaluation. **(A)** Representative photomicrograph of dLight expression (green) in nucleus accumbens (NAc) with DAPI (blue). Fiber tract for optic fiber is demarcated by straight dashed line. **(B)** Injection and implant schematic, wildtype mice were injected with dLight1.1 in NAc and optic fibers at same sites. **(C)** Left panel, mean z-scored NAc dopamine timecourse response during CS test. Following aversion, NAc dopamine release was comparable during responses to CSs paired with either saline or LiCl (0 s = CS response). Right panel, responses collapsed across (pre-CS) baseline and CS response. Main effect of cue presentation only, # p<0.001. **(D)** Left panel, time course responses during consumption test, NAc dopamine activity was elevated during retrieval of devalued reward memory as mice made contact with the food pellets (0 s = pellet response). Right panel, responses collapsed across baseline (5 s prior to pellet response) and pellet response. 1111 Time X condition interaction, p<0.01. Post-hoc Bonferroni group differences **p<0.001.

### Modelling mediated learning and dopamine activity using the Successor Representation Model

The observation that VTA dopamine cells appear to encode detailed reinforcement signals necessary for devaluation of reward memories suggests a novel role of dopamine function that cannot be readily accounted for by standard RPE models of dopamine function ^30^. These models have no mechanism by which the value of a reinforcer can update after CS devaluation.

Dopamine neurons may receive “model-based” inputs that confer devaluation sensitivity ^31,32^; however, it remains unclear what computational function RPEs play within a model-based reinforcement learning system. A related hypothesis is that dopamine neurons signal a vector-valued “generalized prediction error” over a collection of features, rather than just rewards ^33^. This hypothesis can account for a wide range of non-classical dopamine responses, including the sensitivity of dopamine neurons to sensory prediction errors ^7,34,35^ and the causal role of dopamine in learning sensory predictions ^10,36^.

We applied the successor representation (SR) model developed by Gardner and colleagues ^33^ to our mediated devaluation experiment (see supplementary materials for details). This model computes a prediction error for each sensory feature and treats the aggregated dopamine signal as a superposition of these errors. To capture optogenetic and chemogenetic perturbations, the model applies a modulation of the prediction errors. Sucrose consumption is modeled as a monotonic function of expected future reward. The model was able to recapitulate key empirical findings reported above, including the observed reduction in sucrose consumption after CS devaluation with LiCl (Fig. 5A), which was accounted by the capacity of the CS to activate a predictive representation of the sucrose US during aversion, linking it to LiCl by error-driven learning. Moreover, consistent with our behavioral findings (Supplementary Fig. 1) and other studies ^15,25^, the SR model predicts that sucrose consumption will be significantly lower in the minimal relative to the extensive trained condition (Fig. 5B). This reflects the acquisition of a stronger association between the US feature with reward after extensive training, partially counteracting the effects of devaluation during aversion. In addition, the model accurately predicted the dopamine manipulations, which include an enhanced devaluation effect after optogenetic stimulation (ChR2), and a reduced devaluation effect after chemogenetic inhibition (hM4Di) (Fig. 5A). These aspects of the model reflect its capacity to encode predictive representation of sensory features, such that stimulating or inhibiting these errors produces directional changes in stimulus-stimulus learning ^37^. Finally, consistent with the dLight findings, the model predicted higher dopamine transmission during the consumption test in the devaluation condition compared to the control condition (Fig. 5C), due to the surprising absence of gastric malaise that elicits a larger prediction error in the LiCl condition.

**Fig. 5.**
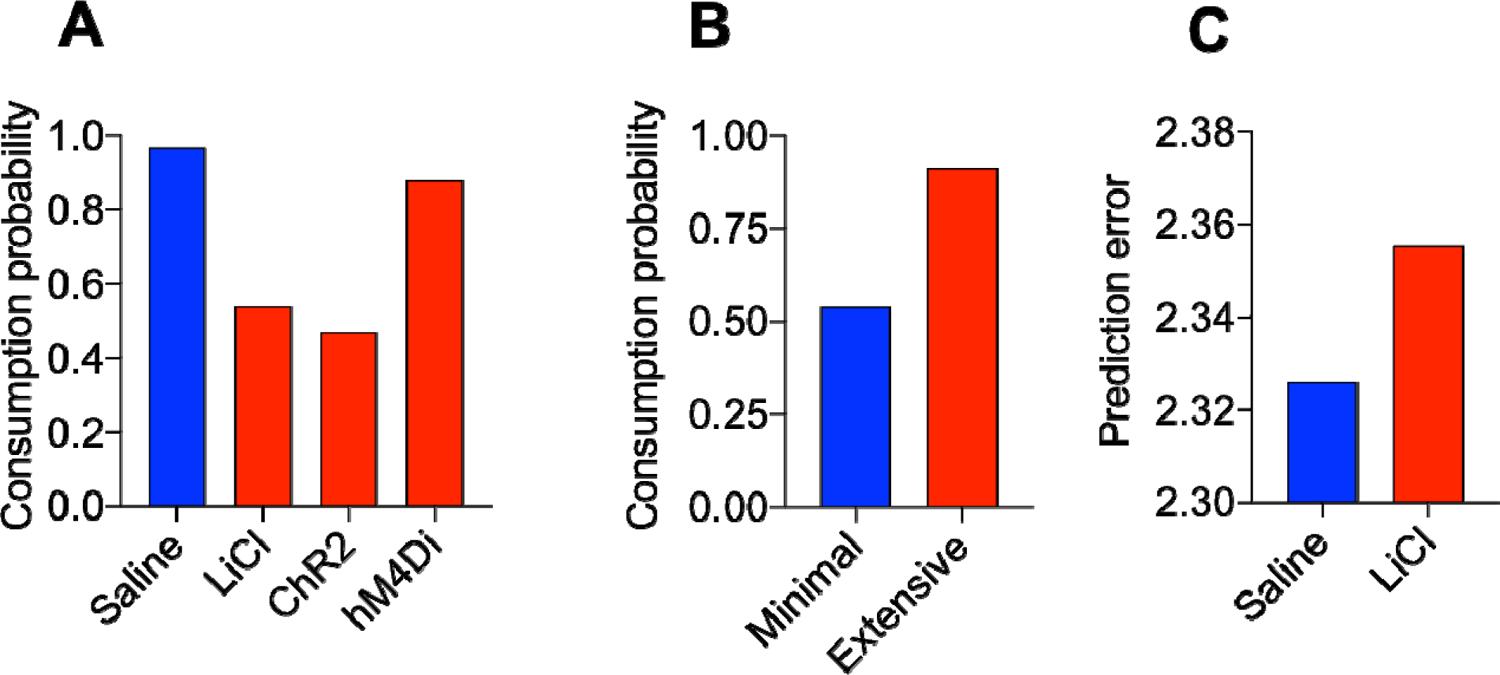
Modelling mediated devaluation through sensory prediction errors. **(A)** Successor Representation model simulation of sensory prediction errors revealed a reduction in sucrose consumption after memory devaluation (Saline vs. LiCl), an augmented devaluation effect following optogenetic stimulation (ChR2), and an attenuated devaluation effect as a result of chemogenetic inhibition (hM4Di). **(B)** The model predicts that mediated devaluation is weaker following extensive Pavlovian training due to the sucrose US feature acquiring a more robust association with reward, which counteracts the effects of CS devaluation. **(C)** The model predicts greater prediction errors during the consumption test in LiCl relative to saline condition.

## Discussion

Our findings reveal a novel function for midbrain dopamine cells through which they gate access to a vast array of detailed features of reinforcement that are typically activated by biologically meaningful stimuli alone. Having identified the parameters underlying mediated devaluation, we demonstrated that mesencephalic cells encode CS-evoked mediated devaluation of sucrose reward. Accordingly, chemogenetic reactivation of these cells led to devaluation due to an associatively mediated reduction in the hedonic taste properties of the sucrose. We also showed that within the VTA, transient activation of dopamine cells during aversion was sufficient to enhance encoding of detailed sensory reinforcement features, which promoted mediated devaluation when mice were subsequently tested in the absence of optogenetic manipulations.

Conversely, chemogenetic inactivation of VTA dopamine neurons prevented perceptual processing of the taste features of the sucrose reward, disrupting the capacity of the CS-LiCl pairings to establish taste-illness associations and elicit a devaluation to the sucrose reward during consumption testing. The retrieval of these reinforcement features also reflected increased dopamine binding in the NAc. These features of dopamine encoding were accurately predicted by the SR model, in which a prediction error for each sensory feature was computed and revealed core features of the dopaminergic manipulations and activity profile of dopamine transients that had been observed during mediated devaluation testing.

These findings stand in stark contrast to traditional accounts ^3,5,38,39^ that restrict dopamine function to encoding RPEs using model-free or cached value signals of future events. These algorithms limit dopamine’s role to encoding value signals, but not specific details about the contents of learning. As such, they are unable to account for why acute dopamine stimulation during aversion would enhance the capacity of CS-LiCl pairings to subsequently augment mediated devaluation to the specific taste features of sucrose reward. Dopamine’s role in this process suggests that in the presence of the CS, stimulation during aversion enhanced further access to detailed sensory (e.g., taste) processing of the associated absent sucrose US, whereas inactivating VTA dopamine neurons at this time prevented retrieval of detailed reinforcement features that were necessary for the generation of mediated devaluation. Moreover, model-free learning systems cannot account for why putative dopamine transients in the NAc would be greater for the food pellet whose memory had been devalued by pairing its CS associate with LiCl. In this latter situation, dopamine activity would be reliant on the transfer of value between cues retrospectively (e.g., food pellet—CS—LiCl), which is not factored into the computational architecture of model-free or cached value reinforcement agents. This heightened activity to the devalued food pellet is all the more challenging given the aversive nature of the phenomenon (i.e., via LiCl) and the prediction by RPE that this should serve to diminish dopamine responses^33^.

More recent findings have questioned the ubiquity of RPE in accounting for dopamine encoding, including that dopamine transients to the NAc do not display RPE in response to encoding positive outcomes (i.e., shock avoidance) ^40^, that dopamine neurons have been shown to fire in response to novelty independent of value ^7^, and appear to encode more specific features of the reward environment including stimulus contingencies ^8^ and detailed information about the characteristics of predicted events ^10^. To account for these findings, investigators have suggested alternative accounts of dopamine function ^41,42^, including that dopamine neurons may receive model-based signals, which allow for computing more detailed associative features of the reward environment ^12^. However, with these algorithms, it is unclear what computational function reward prediction errors play as even though RPE’s are effective in updating value-based estimates of reward and transitions between states in model-based systems, these algorithms do not require prediction errors. Conversely, the SR model assigns prediction errors to sensory signals and treats the overall dopamine signal as a superposition of these errors ^33,43^. By adopting this model, we were able to replicate key features of the mediated devaluation phenomenon. This includes the curious feature that the expression of mediated devaluation generally diminishes as conditioning proceeds (Supplemental Figure 1). Although descriptive accounts have been posited (e.g., extensive training narrows the window of CS-evoked access to detailed US features; ^15,25,44^), mechanistically accounting for this phenomenon has been complicated by the finding that it does not reflect a loss of associability of the CS as it becomes a predictor of the US ^25^. The SR model accounts for weaker mediated devaluation after extensive training because the US feature has a stronger association with reward, and this counteracts the effects of CS devaluation. Moreover, the model makes the prediction that larger RPEs will result from consumption of the devalued food following minimal training, and the devaluation effect should be stronger to the US than the CS following CS-LiCl pairings (Supplemental Figure 8).

It is important to acknowledge the rigorous nature of our approach that combined well-defined associative learning methodology with quantitative measures of ingestive behavior to reveal the nature of the devaluation effect produced by mediated learning. Importantly, changes in sucrose consumption that followed CS-LiCl pairings disrupted the average size of clusters of licks, which reflects a reduction in the palatability of the stimulus ^20,45^. In contrast, measures of licking microstructure that reflect the motivation to engage in sucrose consumption were unaffected by mediated devaluation indicating a selective effect on reducing the hedonic and taste features of the sucrose US. Moreover, consistent with other studies ^14^, directly pairing the CS with LiCl has little impact on subsequent CS processing and is in line with the idea that exteroceptive auditory cues struggle to become associated with illness. Rath er, our findings agree with the account that the CS gated access to the perceptual and sensory features of the US, which became associated with LiCl, leading to devaluation of these reinforcer features. Another important feature of our design is that reductions in sucrose licking did not reflect direct motoric reinforcement following VTA dopamine cell stimulation ^46^, due to the absence of consummatory responses during aversion (i.e., at the time of optical and chemogenetic manipulations).

From a neurobiological perspective, our findings suggest that the encoding of detailed reinforcement signals by VTA dopamine cells is in part relayed to the NAc. The NAc is composed mainly of medium-size spiny neurons (MSNs), which belong to two distinguishable populations of cells based on their expression of dynorphin, substance P and D1Rs, compared to those expressing enkephalin, adenosine A2a receptors, and D2Rs ^47,48^. Interestingly, the capacity of acute stimulation of VTA dopamine neurons to augment mediated devaluation was dependent on D2R activation and there was a tendency for systemic disruption of D2R signaling to impair mediated devaluation. It is tempting to speculate that the circuit underlying dopamine-based encoding of mediated devaluation includes VTA efferents to the NAc in a D2R-dependentr manner. Of course, other target regions from VTA dopamine cells likely play an important role in mediated devaluation, including the basolateral amygdala ^49^, hippocampus ^50^ and other cortical targets such as the insular ^51^ and oribitofrontal cortex ^52^. Future studies targeting the nature of any underlying circuitry encoding detailed reinforcement signals via dopamine should be explored.

Several limitations with our design deserve additional consideration. First, the majority of our studies utilized a single-outcome design, which are not ideally suited to examine sensory-specific encoding ^53^. However, as mentioned above, we have rigorously confirmed that our mediated devaluation effects selectively attenuated the taste and sensory features of the sucrose reward, providing confidence that our manipulations disrupted sensory and not more diffuse features of reinforcement that would have been revealed through other licking (i.e., cluster number) and Pavlovian approach measures. Moreover, we adopted a multiple-outcome design with the fiber photometry experiments and—consistent with the single outcome designs— revealed a pattern of dopamine binding consistent with sensory-specific encoding. Second, our functional manipulations affected all VTA dopamine cells, whereas it is likely that only a subpopulation of these cells underlie mediated devaluation encoding. Future studies employing projection- and cell-specific strategies would be beneficial to distinguish between subsets of dopamine neurons that encode sensory prediction errors. Third, our activity-dependent labelling study include many non-dopaminergic mesencephalic cells. This could reflect GABAergic or glutamatergic involvement ^54,55^, or alternatively leaky expression produced by the manipulation. On a related note, the TH-Cre mouse line employed in the optogenetic and chemogenetic studies is known to suffer from ectopic expression of Cre-recombinase ^56^, and although other models exist (e.g., DAT-Cre), they to suffer from host of other confounding physiological effects that include sex-dependent alterations in associative learning, as well as DAT expression in the striatum ^57^. In addition, our dLight studies were performed in wild-type B6 mice and produced a pattern of responding consistent with the transgenic studies. Finally, unfortunately pellet retrieval times were not recorded in the dLight studies, preventing analyses to determine whether trial-by-trial variations in dopamine binding predicted pellet retrieval.

These limitations aside, our findings provide novel insight into the role of dopamine in reinforcement encoding suggesting a pivotal role in mediated devaluation. To achieve this, a subset of dopamine cells are required to undergo computational processes that are more elaborate than traditional models of dopamine function predict and may in part reflect dopamine’s role in sensory prediction error ^12,33^. Future studies targeting the nature of any underlying circuitry encoding these detailed reinforcement signals via dopamine should include the extent to which our findings relate to proposed roles in model-based encoding ^12^, or signaling of surprise from afferent sensory systems ^58^ that are relayed to higher-order cortical sites ^32^. Alternatively, given our approach intersects appetition and aversion, it is possible that we are engaging a heterogeneous population of dopamine cells that include those that respond to high intensity sensory stimuli and are aversive when stimulated ^59^. In addition to uncovering novel mechanisms of dopamine action in learning, our studies are relevant to neuropsychiatric endophenotypes of reality testing ^60^, and could be leveraged as a tool to attenuate ingestive behavior ^61^ and substance abuse ^62^ through dampening, via mesencephalic DA manipulations, the memories associated with reward.

## Supporting information

Supplemental Methods, Data and Equations

