## Supplemental Methods, Data and Equations for "Devaluing memories of reward: A case for dopamine"

**Supplemental Information**

**Materials and Methods**

**Subjects**

All mice were initially group housed prior to surgery and thereafter single house for the duration of the study. The cfos-htTA mice were originally obtained from Jackson Labs (Strain # 018306) and contained two co-injected transgenes, cfos-tTA and cfos-shEGFP. The expression of tetracycline transcriptional activator (tTA) and green fluorescent protein (shEGFP) directed to activated neurons by the c-fos promoter. Mice were bred for a minimum of two generations with wild-type C57BLJ mice (Jackson Laboratory). cfos-htTA mice (n=5♂, 8♀) were 8-12 weeks at the time of surgery and maintained on a diet containing 40 mg/kg doxycycline for two weeks prior to the start of surgery. For the optogenetic (n=33 ♂, 25♀) and chemogenetic (n=18♂, 18♀) studies, TH-Cre mice expressing Cre recombinase under the control of the tyrosine hydroxylase promoter (TH-Cre) (Jackson Laboratory, Strain #008601) were used. TH-Cre were bred up to four generations out with wild-type C57BLJ mice (Jackson Laboratory). TH-Cre mice were 8-12 weeks at the time of surgery and subsequently single-housed. For the dLight studies (n=♂4,♀2) wild-type C57BLJ mice (Jackson Laboratory) were 10-12 weeks at the time of surgery and subsequently single-housed.

**Virus constructs**

The pAAV-PTRE-tight-hM3Dq-mCherry genome was a gift from William Wisden^63^ (Addgene plasmid # 66795 ; http://n2t.net/addgene:66795 ; RRID:Addgene_66795). The genome was packaged using the triple transfection method and viral particles were isolated from media^64^ and cells, and purified using an iodixanol gradient. Titering was performed using dot-blot as described (*41*) (1.9x10^13^ vector genomes (vg)/ml). ChR2 (AAV5-Ef1α-DIO ChR2-eYFP) and control eYFP (AAV5-Ef1α-DIO-eYFP) were obtained from Vector Labs (Vector Biolabs, Malvern, PA). DREADD (AAV8- hSyn-DIO-hM4D(Gi)-mCherry) and dLight 1.1 (AAV5-CAG-dLight1.1) were obtained from Addgene (Addgene, Watertown, Massachusetts).

**Stereotaxic surgery**

For the activity-dependent labelling study (Experiment 1), at 12 wks of age, cfos-htTA mice were anaesthetized using 5% isoflurane and 1mg/kg buprenorphine, placed in a stereotaxic apparatus and virally infused bilaterally with 0.25 μl of a tet-responsive adenovirus-associated virus pAAV-PTRE-tight-hM3Dq-mCherry expressing hM3Dq at the level of the VTA (AP -3.08, ML +/-0.6, DV -4.5).

To examine the sufficiency of VTA dopamine cells in devaluing memories of food reward (Experiment 2), TH-Cre mice received 0.25μl of a Cre-dependent adenovirus-associated virus expressing channel-rhodopsin (AAV5-Ef1α-DIO ChR2-eYFP; n = 19♂, 16♀) or control eYFP (AAV5-Ef1α-DIO-eYFP; n = 14♂, 9♀) (Vector Biolabs, Malvern, PA) unilaterally at the level of the VTA (AP -3.08, ML +/-0.6, DV -4.5) in a manner counterbalanced for hemisphere. On infection and subsequent recombination in a Cre-expressing cell, the ChR2 version of the virus leads to the expression of modified Na+ channels at the level of the cell membrane that elicit action potentials in the presence of 473nm wavelength light. Viral infusions lacking the ChR2 sequence simply carried a generic eYFP reporter as means of controlling for non-specific effects (e.g. behavior changes due to neural inflammation, tissue damage, etc). Following viral infusions, optic fiber cannulae (200 μm core, 4.1mm; Thorlabs, Newton, NJ) were implanted dorsal to the injection site (AP -3.08, ML +/-0.6, DV - 4.0) and affixed with dental acrylic (Lang Dental Manufacturing Co, Wheeling, IL).

To determine the necessity of VTA dopamine cells (Experiment 3), TH-Cre mice received 0.25μl of a Cre-dependent inhibitory DREADD virus (AAV8- hSyn-DIO-hM4D(Gi)-mCherry; n = 12♂, 12♀) (Addgene) or control eYFP (n = 12♂, 11♀) bilaterally injected into the VTA (AP -3.08, ML +/-0.6, DV -4.5).

For these surgeries, viral injections were performed under isoflurane anesthesia with induction at 5% saturation and maintenance at 4.0% saturation which was dropped by ~1% every 10 minutes until 1.0% maintenance was reached for the remainder of each procedure. Mice were restrained using a Stoelting Brand stereotax (Stoelting Co., Wood Dale, IL). Under aseptic conditions, a dental drill was utilized to drill through the skull at the A.P. and M.L. coordinates before a Hamilton syringe (Hamilton Corp., Reno, NV) could be inserted through the drill site to reach the D.V. coordinates at which point the relevant virus could be infused into the VTA and retained in that position for 5 minutes, after which the syringe was slowly retracted and, if necessary, the process repeated on the contralateral side. The incision site was closed using surgical staples and protected with a layer of triple antibiotic ointment.

To examine *in-vivo* dopamine activity activity in the nucleus accumbens using dLight1.1, mice were deeply anesthetized by inhalation of 2% isoflurane and placed on a Kopf stereotaxic apparatus (Tujunga, CA). Following standard disinfection procedure, the scalp was removed to expose the skull and a small hole was drilled into the skull unilaterally at defined positions to target NAc (A/P: 1.2 mm, M/L: -1.3 mm, D/V: -4.1, -4.5 mm relative to bregma). A pulled-glass pipette with a 20-um tip diameter was inserted into the brain, and the virus was injected by an air pressure system. A picospritzer was used to control injection speed at 25 nl per min and the pipette was withdrawn 5 min after injection. For fiber photometry experiments, 200 µl of AAV5-CAG-dLight1.1 (AddGene #111067-AAV5; titer 7 x 1012 genome copies per ml) was infused. In addition, optical fibers were implanted during the same surgery. Thus, after viral injection, a metal ferrule optic fiber (400-µm diameter core; BFH37-400 Multimode; NA 0.37; ThorLabs) was implanted unilaterally over NAc (A/P: 1.2 mm, M/L: -1.3 mm, D/V: -4.1 mm). Fibers were fixed to the skull using dental acrylic; after the completion of the experiments, mice were sacrificed and the locations of optic fiber tips were identified based on the coordinates of Franklin and Paxinos^65^.

**Drug treatment**

To activate the excitatory (Experiment 1) and inhibitory DREADD (Experiment 3), clozapine-N-oxide (CNO; NIDA Drug Supply Program) powered CNO was diluted in 10% (2-Hydroxypropyl)-β-cyclodextrin in 0.2M sterile phosphate buffered solution (PBS). Mice received intraperitoneal injections of CNO (0.3 mg/kg) 15 mins prior to the memory retrieval and aversion phase. To examine whether the faciliatory effect of optogenetic stimulation of VTA dopamine cells (Experiment 2) during memory retrieval requires intact D2R signaling, mice received a 0.1mg/kg intraperitoneal injection of haloperidol (MilliporeSigma, Burlington, MA) dissolved in a 10% Tween 80 (MilliporeSigma) and 90% sterile saline vehicle. This dose was chosen as it was previously shown to disrupt representation mediated responding without influencing motoric actions^35^.

**Optogenetic stimulation**

For optogenetic stimulation (Experiment 2), 473nm blue light was delivered via a fiber coupled laser source (ThorLabs, Newton, NJ) that was attached to a waveform generator (Agilent Technologies, Santa Clara, CA) integrated into the Med Associates apparatus. For the behavioral studies, prior to each session, the light intensity was tested and calibrated using a high sensitivity power meter (ThorLabs) to emit ∼20 mW at the tip of the 200 µm optical fiber, which was subsequently attached to the ferrule tip of the mouse. Laser stimulation occurred during cue-evoked memory retrieval during the latter 5 s of each CS presentation; the period of time that in training resulted in the delivery of sucrose reward. Mice received 1 s of optogenetic stimulation for (5 ms pulses at 20Hz). For the in-vitro electrophysiology studies (see slice recording, below), slices were exposed to light pulses of 10 Hz and 25 Hz.

**Slice recording**

Tyrosine hydroxylase-positive mice (3-7 months old) were deeply anesthetized with 3% isoflurane and intracardially perfused with cold (~2°C), oxygenated (95% O_2_-5% CO_2_) slicing solution containing (in mM): 2.5 KCl, 1.25 NaH_2_PO_4_, 10.0 MgSO_4_, 0.5 CaCl_2_, 26.0 NaHCO_3_, 11.0 glucose, and 234.0 sucrose, then decapitated. The brains were quickly removed and coronal brain slices (300 µm thickness) containing the VTA were cut using a vibrating tissue slicer (Leica Biosystems, Deer Park, IL) and transferred to a holding chamber containing warmed (~36°C), oxygenated (95% O_2_-5% CO_2_) physiologic solution containing (in mM): 126.0 NaCl, 2.5 KCl, 1.25 NaH_2_PO_4_, 2.0 MgCl_2_, 2.0 CaCl_2_, 26.0 NaHCO_3_, and 10.0 glucose for 30 minutes, after which the holding chamber was maintained at room temperature for the remainder of the experiment. One hour after slicing, individual slices were transferred to a submersion-type recording chamber and superfused (2.5 mL/min) with oxygenated physiologic solution maintained at 32°C.

Recording pipettes were pulled from 1.5-mm outer diameter capillary glass and had tip resistances of 3–5 MΩ when filled with solution containing (in mM): 117.0 K-gluconate, 13.0 KCl, 1.0 MgCl_2_, 0.07 CaCl_2_, 0.1 EGTA, 10.0 HEPES, 2.0 Na_2_–ATP, 0.4 Na-GTP, and 50 μM Alexa Fluor 594. The pH of this solution was adjusted to 7.3 and osmolarity was adjusted to 290 mOsm. The use of this intracellular solution resulted in an 8 mV junction potential that was corrected for in all whole-cell voltage measurements. Data were acquired using a Multiclamp 700B amplifier (Molecular Devices, San Jose, CA) in either current-clamp or voltage-clamp mode. Data were filtered at 4 kHz and digitized at 10 kHz using a Digidata 1440A digitizer (Molecular Devices, San Jose, CA), and collected using pClamp 10 (Molecular Devices, San Jose, CA).

Cell-attached and whole-cell recordings were made from GFP-positive neurons and were visualized using an Olympus BX51WI microscope equipped with Dodt contrast optics and imaged by laser excitation (820 nm) using a two-photon laser-scanning microscopy system (Ultima, Bruker, Billerica, MA) coupled with a Ti:Sapphire laser (Mai Tai HP, MKS-Spectra-Physics, Milpitas, CA). Image stacks and maximum projection images were generated using ImageJ (NIH, Bethesda, MD). Activation of ChR2 was elicited with a small diameter (2.5 µm) single-photon visible laser (473 nm, < 2 mW, 5 ms duration) coupled into the scan head with a photoactivation module and focused over the soma using a second set of galvanometers (Bruker, Billerica, MA).

**dLight recordings**

Beginning 3 weeks after surgery, mice were connected to a fiber optic patch cable. Fiber optic patch cables (0.8m long, 400 μm diameter; Doric Lenses) were firmly attached to the implanted fiber optic cannulae with zirconia sleeves (Doric Lenses). LEDs (Plexon; 473 nm) were set such that a light intensity of <0.1mW entered the brain; light intensity was kept constant across sessions for each mouse. Emission light was passed through a filter cube (Doric) before being focused onto a sensitive photodetector (2151, Newport). Signals were digitized at 60 Hz using PyPhotometry *(43)*, which allows for pulsed delivery of light, minimizing the amount of bleaching over the course of recordings.

To address photobleaching over the course of the recording period, the photometry signal was corrected by subtracting a double exponential fit then adding back the mean of the trace. Signals were then smoothed with a 120 ms sliding window and background subtracted. The fluorescence signal was converted to ΔF/F ((F – F0)/F0; where F0 was calculated as the 10th percentile of the entire fluorescence trace). These traces were then z-scored using the MATLAB (Mathworks, Natick, MA) *zscore* function to facilitate comparisons across days and mice.

**Immunohistochemistry**

Mice were deeply anaesthetized by way of intraperitoneal injection of sodium pentobarbital, then sacrificed via exsanguination during transcardial perfusion with 4% paraformeldahyde (Sigma-Aldrich, St. Louis, MO). Brains were extracted and placed in a 10% sucrose with 4% paraformeldahyde solution for 24 hours at 4° C. Afterward, brains were sliced using a freezing microtome at 30 μm and moved through six, 8 minute washes in 0.1 M phosphate buffered saline (PBS). Next, slices were placed in a solution consisting of 3% normal donkey serum (NDS; Catalog# 017-000-121; Jackson Immunoresearch, West Grove, PA) and 10% Triton-x (MilliporeSigma, Burlington, MA) in PBS for 1 hour, then 24 hours in a solution of 3% NDS, 10% Triton-x, and rabbit-anti-TH primary (Catalog# P21962; MilliporeSigma) at 1:1000 concentration in PBS. The next day, slices were washed six times for 8 minutes each in PBS, then placed in a solution consisting of 3% NDS, 10% Triton-x; and for hM4Di-treated tissue, Alexa-Fluor donkey-anti-rabbit-488 secondary (Catalog# A21206; Invitrogen, Carlsbad, CA), or for ChR2 or eYFP-treated tissue, Alexa-Fluor donkey-anti-rabbit-568 secondary (Catalog# A10042; Invitrogen) at 1:1000 concentration in PBS for 24 hours. Slices were then flow-mounted using 0.1 M phosphate buffer (PB) onto microscope slides and left to dry for 24 hours. Finally, slides were treated with Prolong Gold Antifade Mountant with DAPI (Thermo Fisher Scientific, Waltham, MA) and left to cure for an additional 24 hours prior to imaging and quantification.

**Quantification of hM4Di and ChR2**

Fluorescent images were captured on an Olympus BX51 epi-fluorescent microscope equipped with DAPI, FITC and CY3 filters and connected to an IBM-compatible Windows 10 computer with Neurolucida imaging software (MBF Bioscience, Williston, VT). Relevant brain sections were localized under DAPI using landmarks from Franklin and Paxinos *(42)* and include ventral mesencephalic target sites at bregma: -2.54, -2.7, -2.8, -2.92, -3.08, -3.16 and -3.2. Images were taken at each site and two raters who were unaware of the viral conditions quantified the extent of hM4Di, ChR2 or eYFP expression at each coronal plane. Spread was defined using the following criteria: -/- = no expression, +/- minimal expression, + moderate expression, +/+ extensive expression. In addition, fluorescence expression was traced for each mouse and layers were collapsed across subjects within each coronal plane producing ‘heat maps’, in which lighter and darker shading respectively indicated minimal and maximal expression. At this time, following confirmation of consistency between raters, mice were excluded based on insufficient expression of eYFP (n = 1♂, 1♀), ChR2 (n = 2♂, 2♀) and hM4Di (n = 1♂, 3♀).

**Behavioral Studies**

**Apparatus**

For Experiments 1-3, behavioral procedures were carried out in eight identical, sound-attenuating conditioning chambers featuring steel rod floors at 0.5 cm spacing, translucent polycarbonate walls, and measuring 24 x 20 x 18 cm (Med Associates, St. Albans, VT). Within each conditioning chamber, a magazine area on the right side contained a cut-out food well, into which 50 µl of 0.2 M sucrose reward was delivered. For consumption testing, 0.2 M sucrose was delivered and controlled by the rate of licking by each mouse such every10 licks ∼10-12 µl of solution would automatically be replenished into the food well via syringe pumps controlled by Med-PC software (Med Associates) on an IBM-compatible computer running Windows XP (Microsoft, Redmond, WA). This was accomplished by fiber optic cabling that projected an infrared beam across the fluid meniscus such that each lick would break the beam and allow for timestamped recording of individual licks. Chamber pans and floors were washed with 70% EtOH between male and female subject runs. Conditioning chambers were located in an enclosed, darkened room, illuminated only by red light. For photometry studies, mice were placed in a modified homecage with a FED3 automated feeding device with standard pellets (Dustless Precision Pellets (20 mg); Bio-Serv).

General Behavioral Procedures

*Food cup training*: Prior to behavioral testing, mice were food restricted to 90% of their baseline weight by limiting food access to a single daily portion of lab chow. Mice underwent one day of food cup training in which 50 μl of 0.2 M sucrose was freely available at the start of session. Once initial licking occurred, 16 sucrose deliveries occurred, with each delivery occurring under a random time 120 s schedule. At the start of each trial, a new 50 μl bolus of 0.2 M sucrose was delivered congruently with magazine clicker presentation and made available for 10 s, upon which time it was vacuumed off. Mice were moved on to initial conditioning based on having met the criterion of at least 10 s spent licking while the reinforcer was available.

*Pavlovian Conditioning:* To initially reveal the behavioral parameters for mediated devaluation, (group minimal) mice were split along the lines of either to be paired with LiCl (n = 8) or No-LiCl (n = 8) and began four days of conditioning. Each training session lasted approximately 30 min, during which time they received 4 pseudo-randomly distributed presentations of the 10 s CS with a variable ITI of 450 s. 5 s into each CS, a delivery of a 50 μl 0.2 M liquid sucrose reward (US) occurred. This limited training of 16 CS-US pairings was selected based on pilot studies aimed at setting the conditions for early-stage learning, a period in which representation mediated learning is thought to most readily occur as in *(13)*. To confirm the transient nature of mediated devaluation, an additional group of mice (group extensive) were similarly split along whether they were to be paired with LiCl (n = 8) or No-LiCl (n = 8), however in each session mice received 16 pseudo-randomly distributed trials with the CS and US. This more extensive training exposure with 64 CS-US pairings was expected to prevent subsequent mediated devaluation *(15).* To examine whether optogenetic stimulation during aversion could reignite mediated devaluation in extensively-trained mice, a separate group of eYFP (n = 9) and ChR2 (n = 7).

*Aversion:* After four days of training, mice underwent a fifth day in which they were placed back in the conditioning chamber and allowed to habituate in the absence of cues for 6 min, this was done so as to minimize potential confounding associations as a result of handling stress and to ensure that mice were paying attention to the cue. Following habituation, the CS was played once every 30 s for 5 min in the absence of US delivery, with the goal of rapidly triggering a substitutive CS-evoked representation of the sucrose US. On completion of the session, in a manner counterbalanced for performance during the final two training sessions (i.e., entries/min in the magazine), half of the mice from each group immediately received a 0.6 M intraperitoneal injection of LiCl at 0.15mg/kg before being returned to their home cages. Mice were not fed for 3 hrs post-LiCl to avoid any association between chow and illness. At this stage, cfos-htTA mice (*n*=12) were injected with Dox (66 mg/kg in 10ml/kg) and placed onto a diet containing 40 mg/kg Dox to further prevent labelling. A separate cohort of cfos-htTA mice were euthanized following presentation of either the minimal (n=4) or extensive (n=4) CS and brains processed for cell counting.

*Cue testing:* In order to determine whether the CS entered into direct associations with illness, mice were placed back in their original training context and given four presentations of the CS in the absence of US, entries in magazine during CS presentation was recorded. Mice received four 10 s CS presentations separated by a 450 s fixed ITI.

*Consumption testing:* Mice were placed back in the conditioning chamber for 5 min and given free access to the US in the absence of cues. Licks were recorded for analysis of microstructure with the expectation that a RMTA should occur in the form of decreased average cluster size (the amount of licks contained within a 500 ms pause criterion), a measure known to reflect stimulus palatability.

*dLight studies*: All experiments were conducted as within-subject tests, with mice tested with both flavors of pellets. Only one recording session was conducted on a given day. Animals were food-restricted to 85-90% of their ad lib fed body weight prior to starting behavioral experiments and maintained on a food-restricted diet for the duration of the recording period.

*Habituation to FED3.* To reduce neophobic responses, animals were allowed one 30 min session to acclimate to the recording chamber and freely feed from the in-house FED3 with standard pellets (Dustless Precision Pellets (20 mg); Bio-Serv).

*Habituation to reward.* Animals were allowed two 30 min sessions freely feed from the in-house FED3 with either a banana flavored pellet or berry flavored pellet (Bio-Serv). Animals did not show an inherent preference for either pellet flavor.

*Pavlovian conditioning.* Animals were then trained across 8 alternating 40 min sessions to associate a 1 kHz tone with delivery of the banana flavored pellet and a 2.8 kHz tone with delivery of the berry flavored pellet. ~11 tones were played per session for 10 s, with pellet delivery occurring at 5 s after tone onset, and a variable 3-4 min ITI (using custom Arduino code). After conditioning, mice were provided with an injection of saline.

*Mediated aversion.* During a 10 min session, animals were presented with 10 trials of the 1 kHz tone, with a fixed 30 s ITI. At the end of this session, animals were administered an injection of LiCl. On the following day, this was repeated with the 2.8 kHz tone, after which animals received an injection of saline.

*Reward test.* Animals were allowed one 30 min session to freely feed from the FED3 with a single flavor of pellet and a fixed 1 min ITI. The following day, this was repeated with the other flavor of pellet.

*CS test.* During a 40 min session, animals were presented with 11 trials of the 1 kHz tone, with a 3-4 minute ITI. The following day, this was repeated with the 2.8 kHz tone.

**Data analysis**

Data were subjected to repeated measures analysis of variance (ANOVA). All significant two-way interactions were followed up by repeated measures ANOVA and simple main effects analyses to examine the nature of these interactions. Post-hoc comparisons were analyzed using Bonferroni tests. The α level for significance was .05 and all analyses were conducted using Statistica (Statsoft, Tulsa, OK). In addition, Cohen’s *d* effect sizes measures were calculated (effect size interpretation: small, *d* =.20; medium, *d* =.50; large, *d* =.80).

**Computational modeling**

We adapted the model developed by Gardner et al. (2018)^33^, which we summarize here. The mediated learning paradigm was analyzed into a set of states that are traversed sequentially. Each state *s* was represented by a feature vector f(*s*) = [f_1_(*s*),…, f_D_(*s*)]. We defined 4 features: a constant context feature, a CS feature, a US (sucrose) feature, and a "physiological state" feature (-1 for LiCL, 0 for saline/baseline).

The model formalizes sensory predictions using the *successor representation* [SR; (*9*, *10*), the expected discounted activation of each feature *j* conditional on state *s_t_* is the state at time *t*:

$$M\left( s_{t},j \right)= E\left[ \sum_{k=0} \gamma^{k}f_{j}{(s}_{k}) \right]$$

where E[] is the expectation operator (which averages its argument over randomness in future state transitions), and $\gamma$ is a discount factor (controlling the effective time horizon of the SR). Because in general the state space is large (or possibly infinite), we use a linear approximation of the SR parametrized by weight matrix *W*:

$$\hat{M}\left( s_{t},j \right)= \sum_{i} f_{i}\left( s \right)W_{ij}$$

The weights can be learned using a form of temporal difference learning, analogous to models of reinforcement learning but generalized to arbitrary features:

$$\Delta W_{ij}= \alpha_{W}\delta_{t}f_{i}(s_{t})$$

where $\alpha_{W}$ is a learning rate, and $\delta_{t}$ is a vector-valued prediction error:

$$\delta_{t}\left( j \right)=f_{j}\left( s_{t} \right)+\gamma\hat{M}\left( s_{t+1},j \right)- \hat{M}\left( s_{t},j \right)$$

We model the total dopamine signal in our experiments as a superposition of these errors:

$$DA_{t}= \sum_{j} \delta_{t}(j)$$

Optogenetic and chemogenetic perturbations were modeled by the following equations:

$${\delta'}_{t}\left( j \right)=\left\{ \begin{aligned} \left( 1+\eta\right)\delta_{t\left( j \right)}, \eta<0 \\ f_{j}\left( s_{t} \right)\eta+\delta_{t}\left( j \right), \eta>0 \end{aligned} \right.$$

where $\eta=1.0$ for excitation and $\eta=-0.8$ for inhibition (see Gardner et al., 2018, for justification of this functional form). Obviously, it is a gross oversimplification to model optogenetic and chemogenetic perturbations in the same way, but for our purposes this simplification was adequate to account for the experimental results.

Value computation was modeled by assuming a linear approximation:

$$\hat{V}\left( s_{t} \right)= \sum_{i} f_{i}\left( s_{t} \right)\sum_{j} U_{j}W_{ij}$$

where *U_j_* is a reward prediction weight for feature *j*, updated by an error-driven learning rule:

$$\Delta U_{j}= \alpha_{U}f_{j}\left( s_{t} \right)[r_{t}- \hat{V}\left( s_{t} \right)]$$

with learning rate $\alpha_{U}$. To model consumption choice in the test phase, we transformed the values into choice probabilities using a sigmoidal transformation $F(\hat{V}\left( s_{t} \right)-\tau)$, where *F* is the standard normal cumulative distribution function, and $\tau$ is a response threshold.

To comply with the empirical observation that negative prediction errors have a smaller dynamic range than positive prediction errors (*11*), we rescaled the negative errors for both *W* and *U* by ¼ (though our results don’t depend strongly on this assumption).

We used the same parameter values as in^33^ : $\gamma=0.95, \alpha_{w}=0.06, \alpha_{U}=0.03$. In addition, we set the response threshold $\tau=1$, but the results are qualitatively unchanged for other choices of threshold. Code for the Gardner model can be obtained at https://github.com/mphgardner/TDSR.

Supplemental Figure 1.

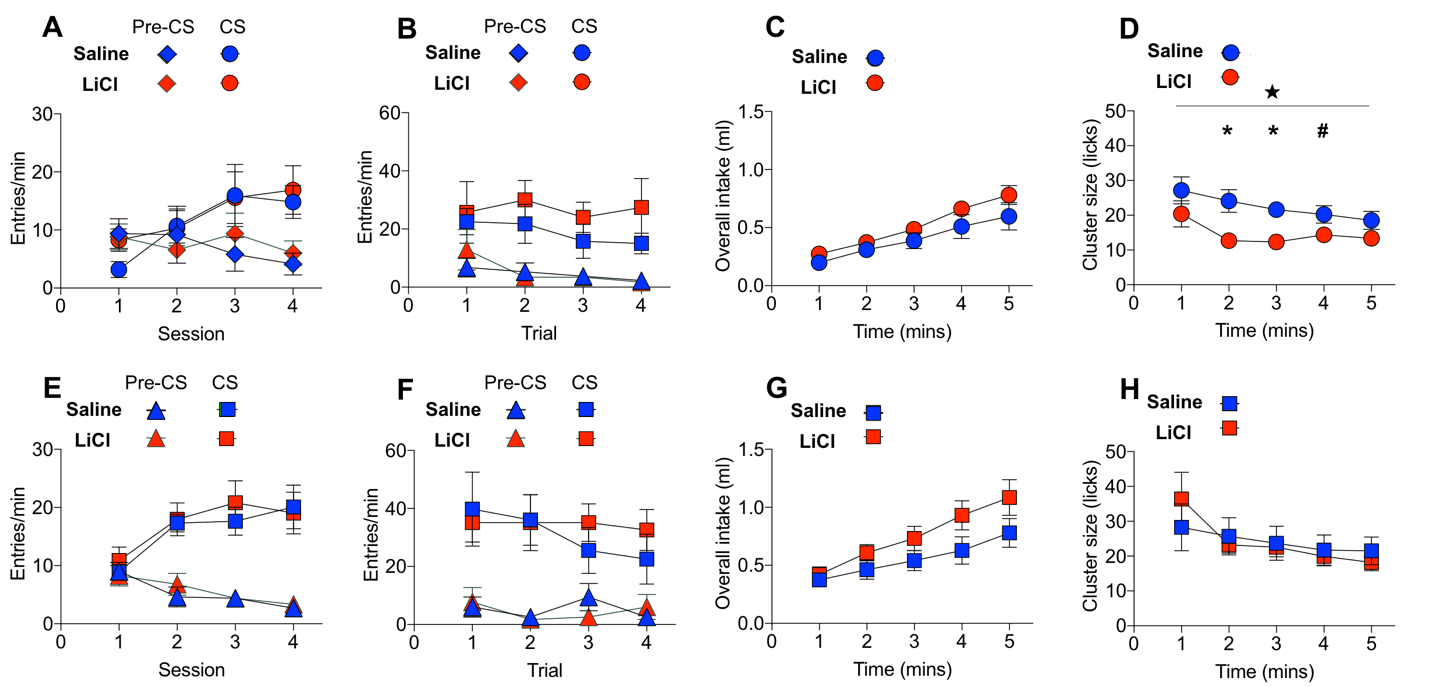

Supplementary Figure 1. Mediated devaluation of sucrose reward reduces its palatability. (A) Mice exposed to minimal Pavlovian training (16 trials) displayed comparable responses to the CSs regardless of the upcoming aversion condition. (B) Following aversion, CS processing remained unchanged and (C) similar overall intake of the associated sucrose reward was also revealed (D) Notably, a significant decrease in palatability of the sucrose reward was accomplished by mediated devaluation of sucrose in LiCl compared to saline mice. Consistent with the idea that CS-evoked access to detailed sensory and perceptual memories of reward narrow with more extensive training, mediated devaluation was transient such that (E) more extensive Pavlovian training (64 trials) (H) eliminated mediated devaluation with no effects on (F) cue or (G) overall intake. ★ main effect of condition, p=0.01, *p’s<0.01, #p<0.05.

Supplemental Figure 2.

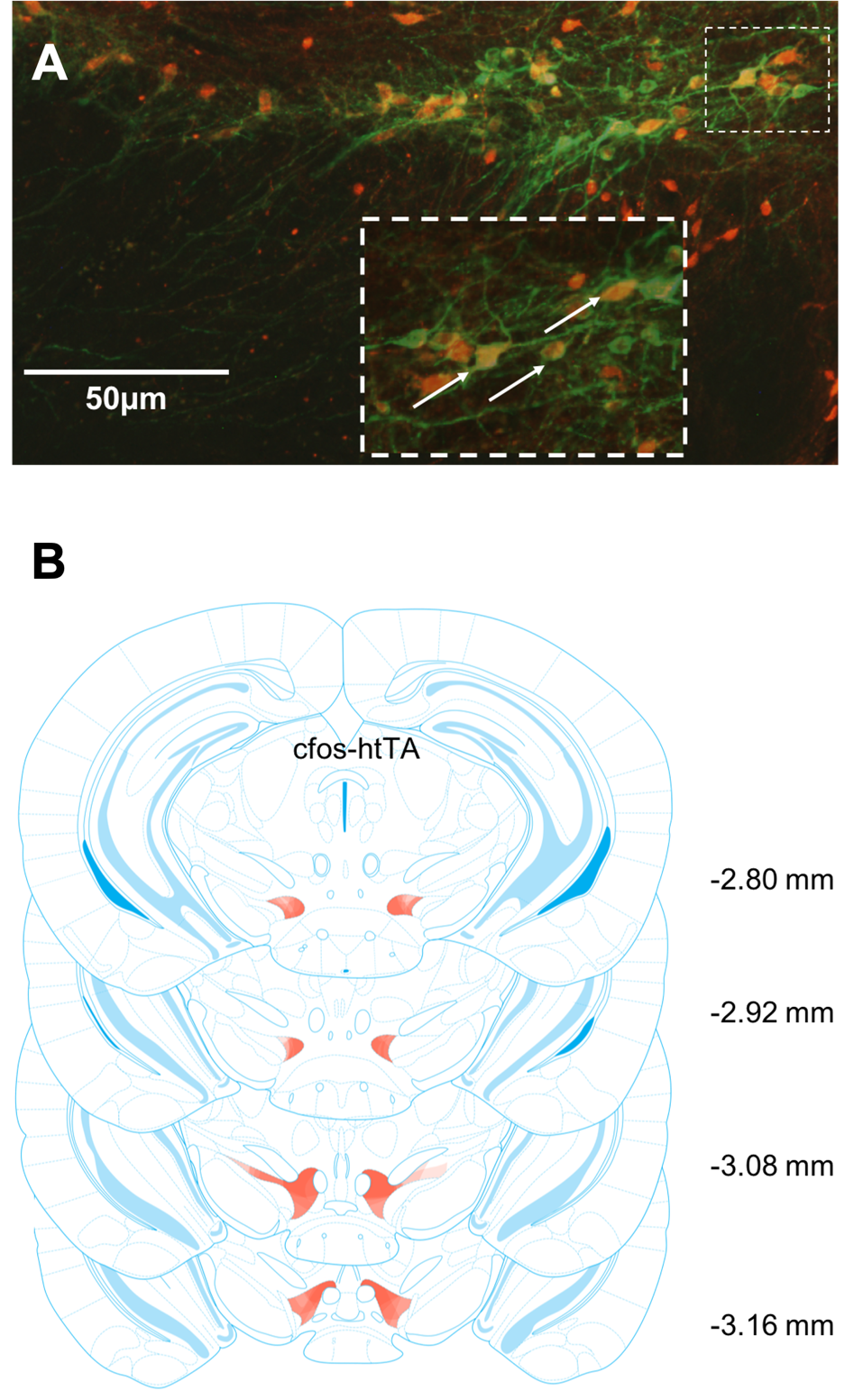

**Supplementary Figure 2.** Expression of activity-dependent labelled hM3Dq expressing cells in ventral tegmental area. **(A)** In cfos-htTA mice, hMe3Dq expression (red) colocalized with tyrosine hydroxylase (TH) positive cells (green) in ventral tegmental area. Arrows indicate somatic expression of hM3Dq. **(B)** Heat maps demarcating the extent of hM3Dq expression in ventral tegmental area. Lighter and darker shading respectively indicate minimal and maximal expression at each coronal plane.

Supplemental Figure 3.

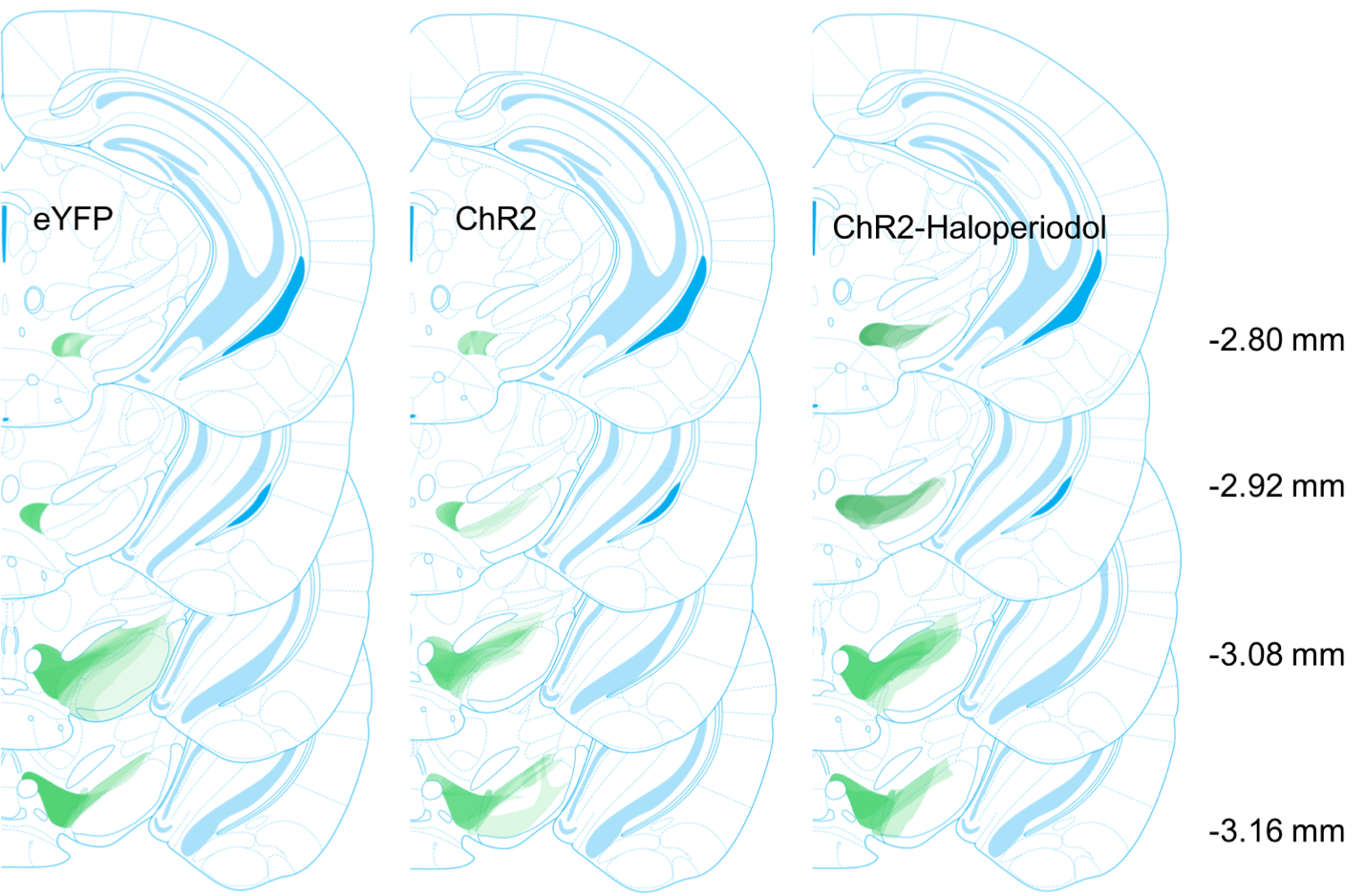

**Supplementary Figure 3. Expression of ChR2 in ventral tegmental area cells.** Heat maps demarcating the extent of ChR2 expression in ventral tegmental area. Lighter and darker shading respectively indicate minimal and maximal expression at each coronal plane.

Supplemental Figure 4.

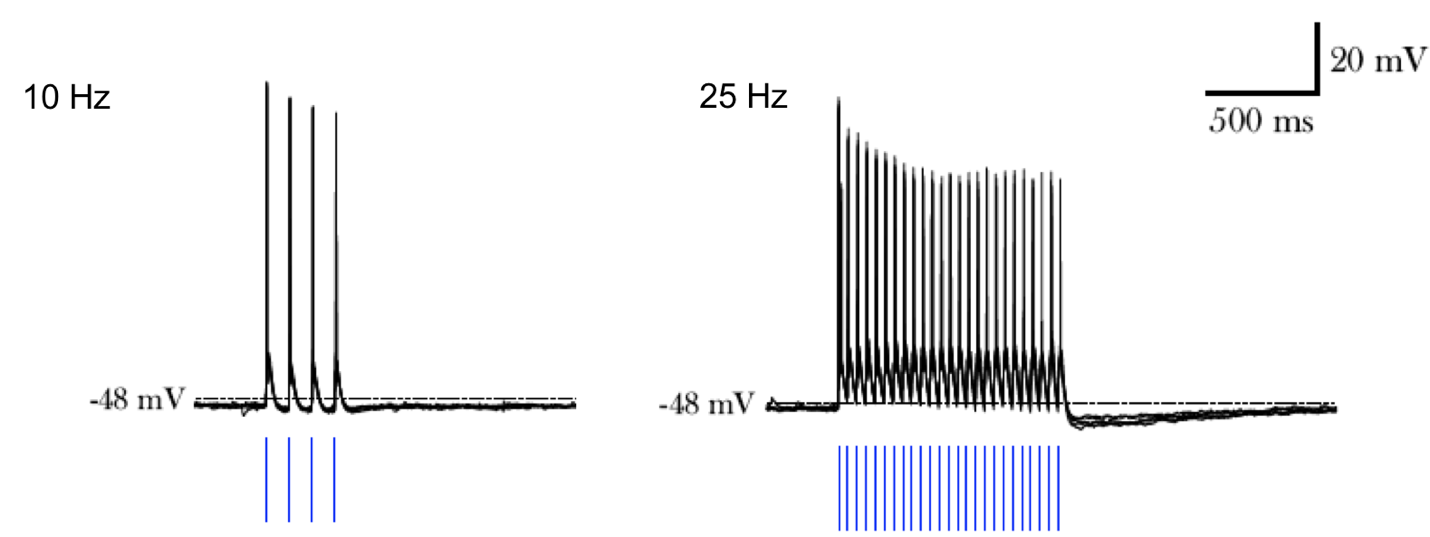

**Supplementary Figure 4.** Brain slice whole-cell voltage responses to pulses of blue light at 10Hz and 25 Hz in ChR2-expressing TH neurons.

Supplemental Figure 5.

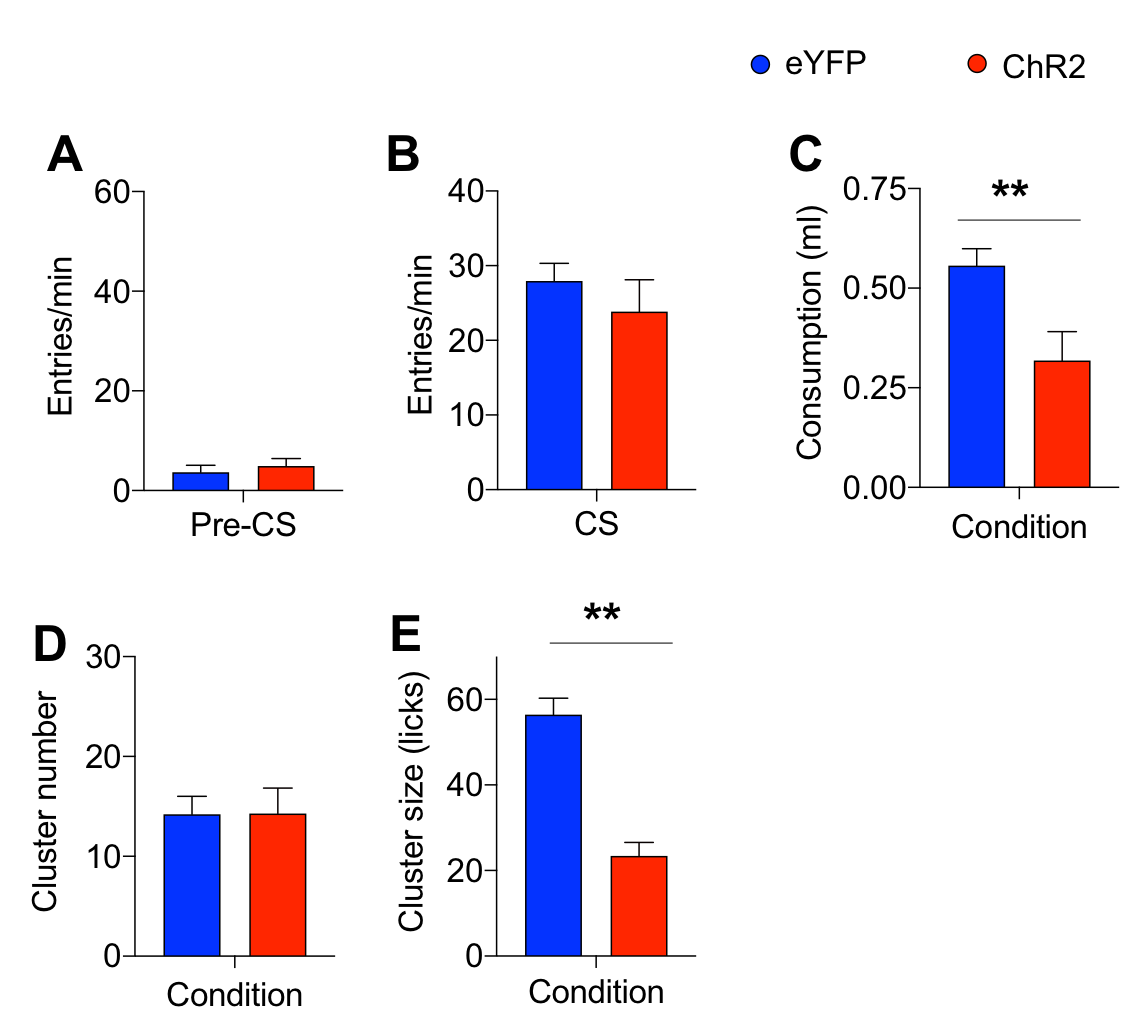

**Supplemental Figure 5.** **Optogenetic stimulation of ventral tegmental area dopamine cells rescues mediated devaluation.** **(A)** pre-CS and **(B)** CS elicited food cup entries were unaffected by prior acute optogenetic activation of dopamine cells that occurred during aversion. **(C)** Laser stimulation during aversion led to a subsequent reduction in overall intake of sucrose in ChR2 mice that were trained under more extensive Pavlovian training conditions that typically limit the expression of mediated devaluation. **(D,E)** Licking microstructure analyses revealed the nature of the devaluation to sucrose reflected a reduction in the perceived palatability of the reward as indicated by **(E)** cluster size, without impacting **(D)** the motivation to initiate sucrose consumption, i.e., burst number. Group differences, ****** p’s=0.01.

Supplemental Figure 6.

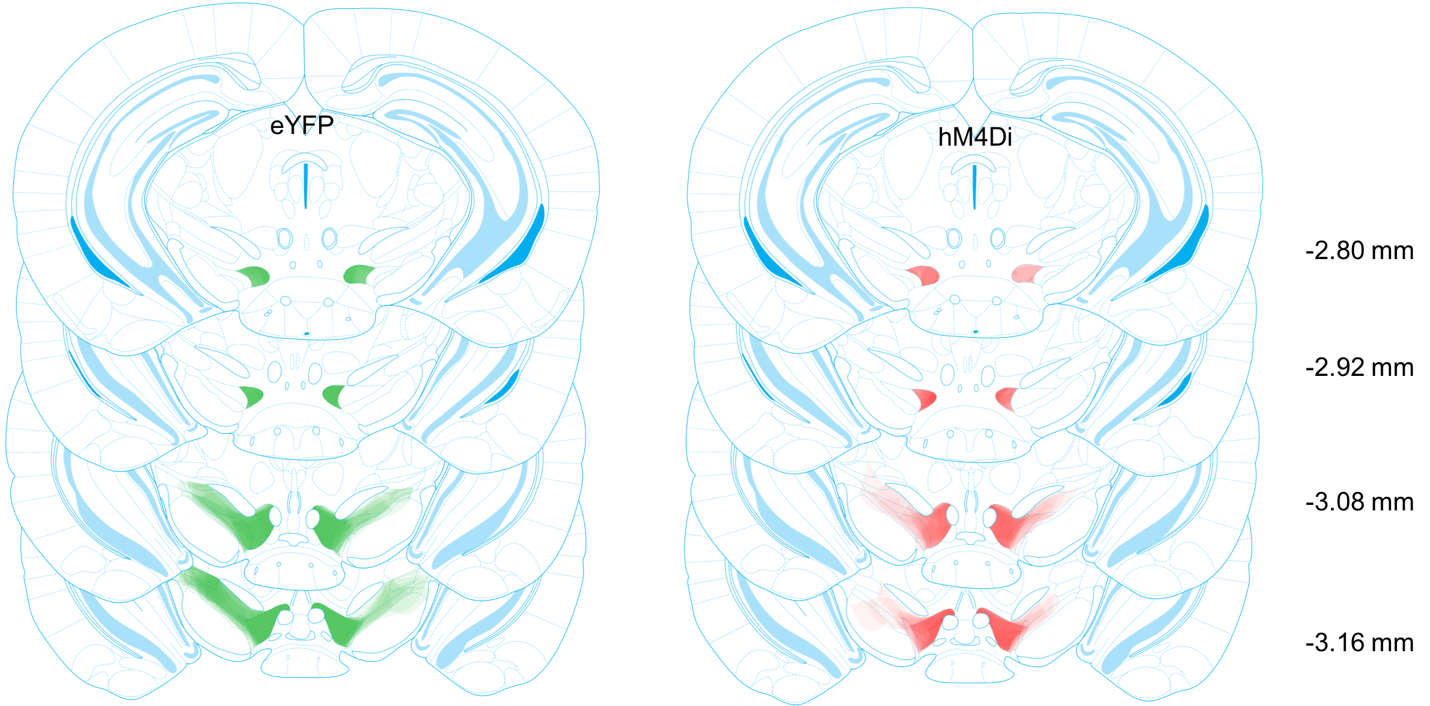

**Supplementary Figure 6. Heat maps showing density of eYFP and hM4Di expression in ventral tegmental area.** In TH-Cre mice, eYFP (green) and hM4Di expression (red) was robustly expressed throughout the ventral mesencephalon. Lighter and darker shading respectively indicate minimal and maximal expression at each coronal plane.

Supplemental Figure 7.

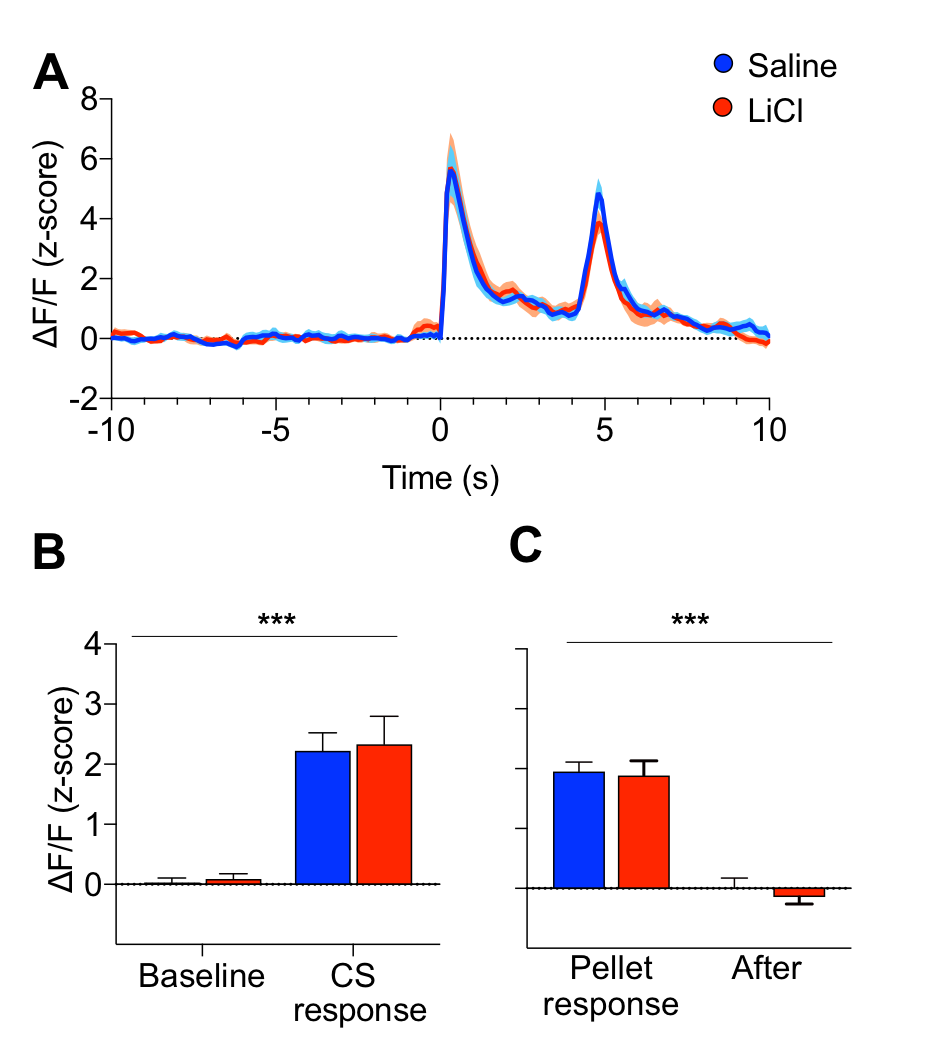

**Supplementary Figure 7.** **Dopamine release in nucleus accumbens during Pavlovian training. (A)** Mean z-scored NAc dopamine timecourse response during the final Pavlovian training session prior to memory devaluation. Data are presented for CS and pellet responses for the CS that would subsequently be paired with saline or LiCl during aversion. **(B)** Responses collapsed across 4 s (pre-CS) baseline and during CS presentation in the final Pavlovian training session. Thus, prior to aversion, NAc dopamine responses were comparable for CS subsequently paired with saline or LiCl. Time period X condition ANOVA (F<1), *** indicates main effect of time period only, p<0.0001. **(C)** Responses collapsed across pellet acquisition (2s) delivery and period following its delivery (2 s). Mice displayed similar NAc dopamine responses to the two distinct food pellets prior to aversion. Time period X condition ANOVA (F<1), *** indicates main effect of time period only, p<0.0001.

Supplementary Figure 8.

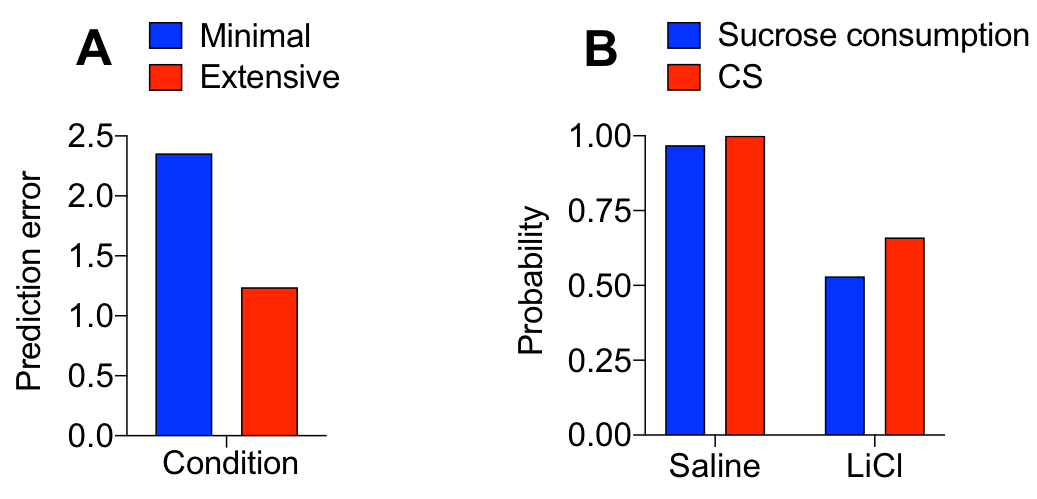

**Supplementary Fig. 8.** **Extensive training produces smaller prediction errors and mediated devaluation to US is stronger. (A)** The Successor Representation model predicts that in the consumption test, prior extensive training produces a stronger expectation for the context feature that is confirmed during the consumption test resulting in a smaller error term. In addition, extensive training produces a stronger expectation of the CS when the US is present, this violation during consumption testing produces a negative prediction error for this feature. **(B)** Stronger devaluation to the US is predicted by the Successor Representation model because although the CS has a negative immediate reward expectation, it also has a relatively strong predictive relationship with the US, which has a positive immediate reward expectation (since it was never paired directly with LiCL)—these two expectations counteract each other. It should be noted that in the generation of this model prediction, we made the limiting assumption that the CS and US conditioned response measures have the same linking assumption to the value estimate, which is unlikely. Nevertheless, with these parameters, the model is able to predict a relatively stronger US devaluation effect.
